# Borrelia PeptideAtlas: A proteome resource of common *Borrelia burgdorferi* isolates for Lyme research

**DOI:** 10.1101/2023.06.16.545244

**Authors:** Panga Jaipal. Reddy, Zhi Sun, Helisa H. Wippel, David Baxter, Kristian Swearingen, David D. Shteynberg, Mukul K. Midha, Melissa J. Caimano, Klemen Strle, Yongwook Choi, Agnes P. Chan, Nicholas J. Schork, Robert L. Moritz

**Author notes:** Equal first authors (Panga Jaipal. Reddy, Zhi Sun, Helisa H. Wippel). Corresponding author: Robert L. Moritz.

## Abstract

Lyme disease, caused by an infection with the spirochete *Borrelia burgdorferi*, is the most common vector-borne disease in North America. *B. burgdorferi* strains harbor extensive genomic and proteomic variability and further comparison is key to understanding the spirochetes infectivity and biological impacts of identified sequence variants. To achieve this goal, both transcript and mass spectrometry (MS)-based proteomics was applied to assemble peptide datasets of laboratory strains B31, MM1, B31-ML23, infective isolates B31-5A4, B31-A3, and 297, and other public datasets, to provide a publicly available Borrelia PeptideAtlas (http://www.peptideatlas.org/builds/borrelia/). Included is information on total proteome, secretome, and membrane proteome of these *B. burgdorferi* strains. Proteomic data collected from 35 different experiment datasets, with a total of 855 mass spectrometry runs, identified 76,936 distinct peptides at a 0.1% peptide false-discovery-rate, which map to 1,221 canonical proteins (924 core canonical and 297 noncore canonical) and covers 86% of the total base B31 proteome. The diverse proteomic information from multiple isolates with credible data presented by the Borrelia PeptideAtlas can be useful to pinpoint potential protein targets which are common to infective isolates and may be key in the infection process.

## Background & Summary

The spirochete *Borrelia burgdorferi* is the causative agent of Lyme disease, the main vector-borne infection in North America, with over 476,000 cases per year between 2010 and 2018 [1, 2]. *B. burgdorferi* is transmitted to humans through the bite of infected nymphal or adult blacklegged ticks, and the untreated infection may cause a multisystem disorder characterized by early and later stage signs and symptoms [3, 4]. The early stage symptoms happen within the first 30 days after the tick bite and may include, among others, fever, joint aches and swollen lymph nodes, and rash (erythema migrans) [3]. When symptoms persist for months after the tick bite, later symptoms could be manifested: facial palsy, arthritis with severe joint pain and swelling, severe headaches, and inflammation of the brain [3]. Treatment is done with the use of antibiotics, and most cases of Lyme disease can be cured within 2 to 4 weeks [5], even though patients may continue to present symptoms for more than 6 months after the treatment ends, condition called Post-Treatment Lyme Disease Syndrome (PTLDS) [6]. Therefore, an early and correct diagnosis of Lyme disease is key to initiate the treatment of infection at its early stages and prevent extreme symptoms. Currently available tests, including the current CDC-recommended 2-tiered testing protocol, are designed to detect antibodies against *B. burgdorferi* in patient blood, which takes several weeks to be produced and can result in a false negative diagnosis [7]. Therefore, the development of alternative diagnostic methodologies, such as next-generation serologic assays [7], which include recombinant proteins and synthetic peptides targeting important factors in the *B. burgdorferi* infection, survival and proliferation mechanisms are needed.

*Borrelia burgdorferi* is an atypical Gram-negative bacteria due to lack of LPS in its cell wall and the presence of immuno-reactive glycolipids, a peptidoglycan layer, and lipoproteins in the outer membrane [8-11] (Figure 1). These lipoproteins play a key role in the infectivity and proliferation of the spirochete in ticks and in mammal hosts [12], and are mostly encoded by the spirochete linear and circular plasmids, besides the single chromosome harbored [13]. Specifically, the *B. burgdorferi* B31 genome sequence revealed the presence of one linear chromosome with 843 genes, and 21 plasmids (12 linear and 9 circular) with 670 genes and 167 pseudogenes [9, 14]. Out of a total of 1,513 genes, 1,291 are predicted as unique protein-coding genes [14]. B31 is the most commonly studied *B. burgdorferi* non-infective laboratory isolate, but an increasing number of infective genotypes have been isolated in North America and around the world, which are isolated from infected ticks or Lyme patients and display different pathogenic and infective patterns [15]. The genetic variability of subtypes of *B. burgdorferi* isolates – e.g., varying number of plasmids encoding for infection-related lipoproteins – may ultimately lead to diverse (i) severity of Lyme symptoms and (ii) spirochetal response to the antibiotic treatment [15]. Hence a proteogenomic approach combining genome sequencing data with transcriptomic and proteomic data from different isolates is a robust strategy to unveil the Borrelia pathogenicity and begin to develop new strategies for more efficient diagnosis and treatment of Lyme disease.

**Figure 1.**
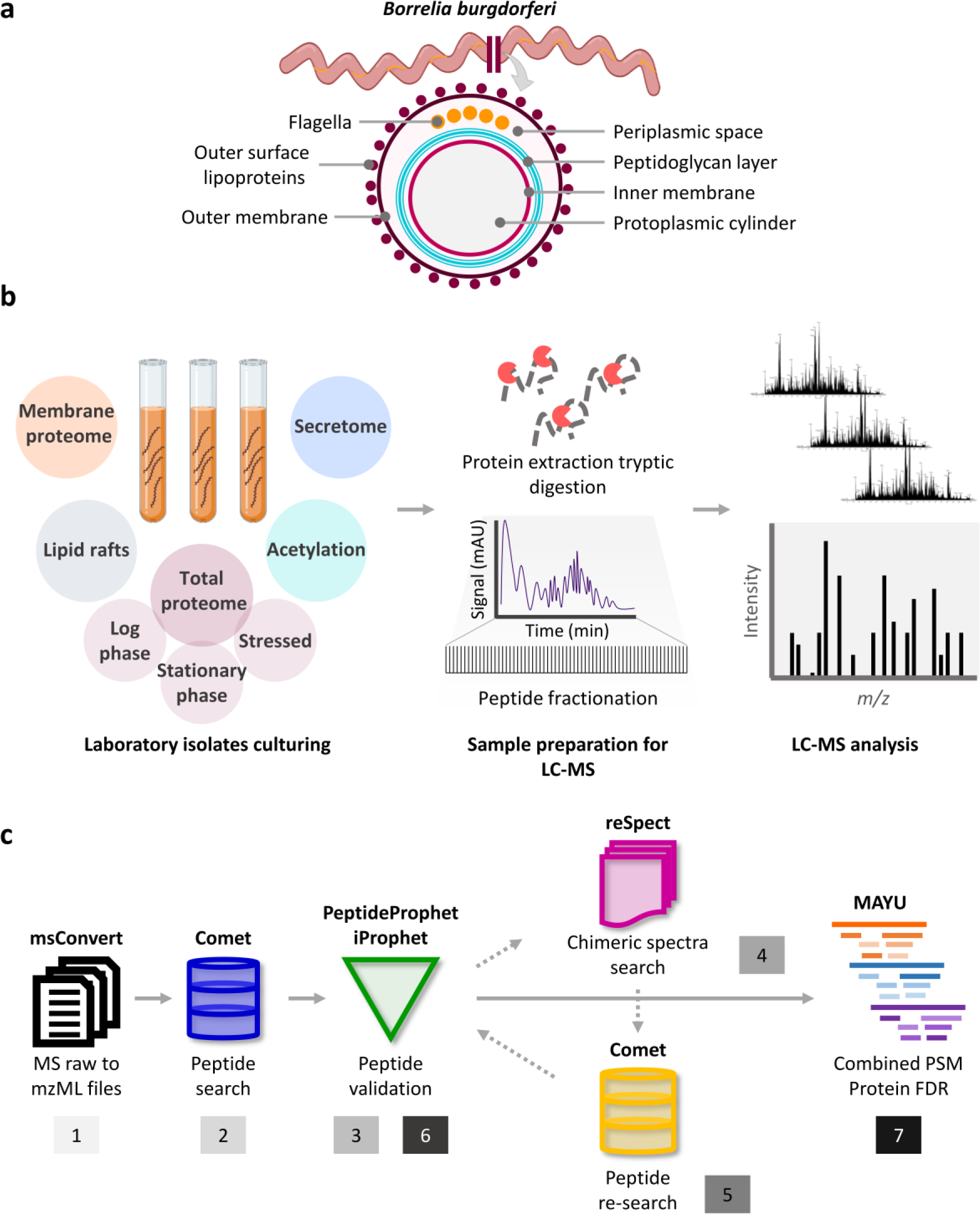
Overview of experimental workflow for the development of the Borrelia PeptideAtlas. (a) Cartoon representing *B. burgdorferi* structure. (b) Experiment workflow. *B. burgdorferi* was cultured in different environmental conditions, including log phase, stationary phase, and stressed conditions for total proteome analysis. Different enrichment assays were applied for the analysis of the secretome, the membrane proteome, lipid rafts, and acetylation. Samples were prepared directly for LC-MS analysis, or alternatively fractionated prior to LC-MS. Further details in Methods. (c) Trans Proteomic Pipeline (TPP) for the Borrelia PeptideAtlas assembly.

Although numerous proteomic reports exist for *B. burgdorferi* isolates [16-23], no information is available as a searchable compendium of public data, and users have to resort to obtaining raw information files and search any, or all, of these data individually. In this study, we include in-house acquired mass spectrometry (MS) and publicly available data to perform a comprehensive proteome analysis of 6 different laboratory *B. burgdorferi* isolates: two commercially available isolates B31 [24] and MM1 [25, 26], the genotype B31-ML23 [27], and 3 infective isolates 297 [28], B31-A3 [18], and B31-5A4 [29]. These datasets include information on the total proteome, secretome, and membrane proteome of the *Borrelia* isolates (Figure 1a&b). The uniform analysis of the total 855 MS runs through the Trans-Proteomic Pipeline (TPP) (Figure 1c) allowed the identification of 76,936 distinct peptide sequences at false discovery rate (FDR) levels less than 0.1%. These unique peptides map to 1,221 canonical proteins among all isolates with a protein-level FDR less than 1% covering 86% of the total B31 proteome coverage. Additionally, for the comparison of protein abundance levels with mRNA levels, we performed transcriptomic analysis of isolates B31, 5A4-B31, and MM1. The complex and detailed proteomic results achieved here were gathered in a public repository called Borrelia PeptideAtlas, with interface made available at http://www.peptideatlas.org/builds/borrelia/. PeptideAtlas is a unique public community resource which contains large scale assembly of mass spectrometry data uniformly processed through the TPP [30, 31]. This repository has data from a wide range of samples, e.g., including human, *Saccharomyces cerevisiae, Drosophila melanogaster* and *Candida albicans* [32-36] among others. The Borrelia PeptideAtlas allows the assessment of protein content of *B. burgdorferi* isolates and compare detectable protein sequences. The continuous update of this repository with expandable data sources for many other *B. burgdorferi* isolates, including clinically relevant isolates, will enable the investigation of the dynamic proteome of this spirochete through its infection stages and their vastly different environments. The diverse proteomic information from multiple infective isolates with credible data presented by the Borrelia PeptideAtlas can be useful to understand the protein complement of each isolate and assist in pinpointing potential protein targets which are common to infective isolates and may be key in the infection process. The Borrelia PeptideAtlas is readily available as an important resource for the Lyme disease research community.

## Methods

### *B. burgdorferi* isolates and spirochete culture

For the in-house performed experiments, 2 common commercially available laboratory isolates of *B. burgdorferi* [B31 (ATCC 35210) [24], MM1 (ATCC 51990) [25, 26]], and the infective isolate B31-5A4 (a clonal isolate of 5A4 that has been passaged through rodents to maintain infectivity) [29] were cultured in BSK-H complete media with 10% rabbit serum, at 34 °C in 5.0% CO_2_ incubator. B31-5A4 was cultured at a low passage to minimize loss of endogenous plasmids. The spirochetes were harvested and collected at mid-log phase (3 to 5 × 10^7^) or stationary phase (3 to 5 × 10^8^) for proteomic analysis. For secretome analysis, mid-log phase cells were harvested, washed and transferred to serum-free media, i.e., BSK-H media without rabbit serum grown for 24 h at 34 °C in 5.0% CO_2_ incubator. The culture was centrifuged at 3,000 rpm for 1 h and collected both the media and the bacteria were collected. The media was used for secretome analysis and bacteria were used for stress proteome analysis.

### Total proteome extraction

*B. burgdorferi* pellets collected from log phase, stationary phase and stressed bacteria (grown in serum-free media for 24 h) were washed with PBS buffer (pH 7.4) four times to remove the media and centrifuged at 300 × *g* for 3 min at each wash. The bacterial pellets were dispersed in lysis buffer of 8 M urea in 100 mM ammonium bicarbonate and protease inhibitor cocktail (CoMplete, (Roche)). The bacterial cell lysis was performed using freeze-thaw cycle followed by sonication (30 s pulse, 20% amplitude, 5 cycles). Cell lysate was centrifuged at 25,000 rpm for 30 min and clear supernatant was collected for total proteome analysis. The protein samples were stored at - 80 °C until used.

### Secretome extraction

*B. burgdorferi* B31 culture at mid-log phase was washed with PBS buffer to remove the media and allow transfer of the bacteria to serum free BSK media (BSK media without rabbit serum) for 24 hrs. The bacteria were collected by centrifugation at 300 × *g* for 3 min, and the media was used for the secretome analysis. To the media, four volumes of chilled acetone were added and precipitated the protein for 30 min at 4 °C. Protein pellets were collected by centrifugation and washed with acetone two more times. Protein pellet was dissolved in lysis buffer (8 M urea in 100 mM ammonium bicarbonate).

### Membrane proteome analysis

*B. burgdorferi* B31 was cultured as above and harvested by centrifugation at 3,000 rpm for 60 min. The bacterial pellets were washed with ice cold PBS buffer (pH 8.0) three times. The bacterial pellets were resuspended in PBS buffer (pH 8.0) with final cell number of 10^8^/mL of buffer. 10 mM Sulfo-NHS-SS-Biotin (Thermo-Fisher Scientific, USA) was prepared according to the manufacture’s guide. The stock solution of Sulfo-NHS-SS-Biotin was added to the bacterial pellets and mixed via pipette. The bacterial pellets were incubated at 4 °C for 60 min for the labeling reaction. Each bacterial pellet was centrifuged at 5,000 rpm for 20 min and supernatant was discarded. Tris buffered saline (TBS, pH 7.4) was added to the bacterial pellets and incubated at room temperature for 15 min and centrifuged at 16,000 rpm for 10 min. *B. burgdorferi* pellets were washed with PBS buffer (pH 7.4) and dispersed in 100 mM Tris buffer (pH 8.0) containing the protease inhibitor cocktail. The cell lysis was performed using freeze-thaw cycles as described above. The *B. burgdorferi* B31 lysate was centrifuged at 25,000 rpm for 30 min and the supernatant was collected for soluble proteome analysis. The resultant protein pellet was washed with 100 mM Tris buffer (pH 8.0) and dissolved in membrane dissolving buffer (8 M urea having protease inhibitor cocktail) and incubated at 4 °C for 30 min with intermediate vortexing. The sample was centrifuged at 25,000 rpm for 30 min and the supernatant was collected for membrane protein analysis. Alternatively, DynaBeads (Dynabeads MyOne Streptavidin T1, Invitrogen) were prepared by adding PBS buffer (pH 7.4). Membrane fractions were transferred to the tubes having beads and incubated for 1 h at 4 °C with end-over-end rotation. Beads were sequestered by a magnet and sequential washing steps were performed as follows: 1 mL per wash and 8 min per wash with solution-I (2% SDS), solution-II (6 M urea, 0.1% SDS, 1 M NaCl and 50 mM Tris pH 8.0), solution-III (4 M urea, 0.1% SDS, 200 mM NaCl, 1 mM EDTA and 50 mM Tris pH 8.0) and solution-IV (0.1% SDS, 50 mM NaCl and 50 mM Tris pH 8). The bound proteins were eluted in 2 × SDS-PAGE sample buffers.

### In-solution digestion and high-pH fractionation

Isolates B31, MM1 and B31-5A4 protein samples (log phase, stationary phase, stressed bacteria, and secretome) were digested with trypsin for proteomic analysis. Briefly, 100 µg of protein from each condition were reduced with 5 mM Tris (2-carboxyethyl) phosphine (TCEP) and alkylated with iodoacetamide. Proteomic-grade modified trypsin (Promega) was added at a 50:1 protein-to-enzyme ratio and incubated at 37 °C overnight. Samples were fractionated using high-pH fractionation, and the remaining samples were analyzed by LC-MS/MS directly. In the first experiment set trypsin digested peptides were reconstituted in 200 mM ammonium formate (pH 10) and fractionated on an Agilent 1200 Series HPLC system. Peptides were loaded on ZORBAX SB-C18 column (4.6 × 150 mm, 5 µm particle size) and fractionated using a linear gradient of 0-100% of B (60% acetonitrile in 20 mM ammonium formate pH 10). A total of 24 fractions were collected over the elution profile and pooled to create 8 disparate fractions, each containing 3 of the original 24, separated by 7 fractions in between, i.e. (1,9,17), (2,10,18), (3,11,19) etc. For B31, B31-5A4 and MM1 protein samples tryptic peptides were fractionated on the Agilent 1200 Series Gradient HPLC system with a flow rate of 100 µL/min of buffer A [0.1% (vol/vol) triethylammonium bicarbonate (TEAB) in water] and 1%/min gradient of buffer B [60% (vol/vol) acetonitrile, 0.1% (vol/vol) TEAB in water], with a Brownlee Aquapore RP-300 column (100 mm × 2.1 mm i.d. from Perkin-Elmer). The total 56 fractions were pooled to 14 final fractions through groupings of 3 disparate fractions to cover the range. These fractions were lyophilized and reconstituted in 0.1% formic acid and 2% acetonitrile for LC-MS/MS analysis.

### SDS-PAGE and in-gel digestion

The biotin labeled proteins eluted in 2 × SDS-PAGE sample buffers were mixed with reducing agent and bromophenol blue (BPB) and resolved on 12% SDS-PAGE gel. The gel was stained with SimplyBlue Safe Stain (Invitrogen, Carlsbad, CA). Each lane of the SDS-PAGE was cut into five bands and processed for in-gel digestion. In brief, the gel pieces were washed with 50 mM ammonium bicarbonate (AmBic) and 2:1 ratio of acetonitrile:AmBic alternatively,three times for five min each to remove the stain. Gel bands were treated with DTT (56 °C for 1 h) and iodoacetamide (20 min in the dark) for reducing and alkylating the cysteine residues. Trypsin (500 ng/µL) along with sufficient 50 mM AmBic was added to each gel band and incubated at 37 °C overnight. Peptide elution was performed by adding 60% of acetonitrile in 0.1% TFA to the bands,vortexing for 10 min and collectint the solution into a fresh tube. The process was repeated two more times with acetonitrile gradient 70% and 80% in 0.1% TFA and pooled to the previous fraction.

### Enrichment of phosphorylated peptides

For the enrichment of phosphorylated peptides of isolates B31-5A4 and MM1, tryptic peptides from log phase pellets (∼5 × 10^7^ cells) were resuspended in 500 µL of loading buffer [80% acetonitrile, 5% trifluoroacetic acid (TFA), 0.1 M glycolic acid], and incubated with 400 µg of MagReSyn Ti-IMAC HP (Resyn Biosciences). Beads were washed 3 times with 500 µL of 80% acetonitrile and 1% TFA, 3 times with 500 µL of 10% acetonitrile and 0.2% TFA, and peptides were eluted with 200 µL of 2% ammonium hydroxide. Samples were cleaned up with a C18 Atlas column (Tecan, USA) and prepared for LC-MS analysis.

### LC-MS/MS analysis

#### Q-Exactive HF

*Borrelia burgdorferi* samples, except B31-Biotin labeled samples, were analyzed either on an EasynLC (Thermo Fisher Scientific) coupled with Q-Exactive HF mass spectrometer (Thermo Fisher Scientific). The purified dried peptides were dissolved in loading buffer (0.1% formic acid (FA) in water) and loaded on to the Acclaim PepMap 100 trap (2 cm long, 75 μm ID, C18 3 μm; Thermo Fisher Scientific). Analytical column (PICOCHIP: 105 cm, 1.9 µm, REPROSIL Pur C-18-AQ, 120 Å, New Objective, USA) with a flow rate of 300 nL/min was used for the separation of the peptides with a linear gradient of 5–35% buffer-B (90% acetonitrile in 0.1% FA) over 120 min. The data acquisition parameters include: mass range 375-1375 *m/z*, MS resolution of 30,000 (at m/z 200), MS/MS resolution of 15,000 (at *m/z* 200), full scan target at 3 × 10^6^, 40 top intense peaks with charge state >2 were selected for fragmentation using HCD with 28% normalized collision energy, dynamic exclusion time of 25 s and profiler mode with positive polarity. Alternatively, B31-Biotin labelled peptides were analyzed using Agilent 1100 nano pump coupled to an LTQ Velos Pro-Orbitrap Elite mass spectrometry (Thermo Scientific, USA). Sample was loaded onto a trap column consisting of a fritted capillary (360 μm o.d., 150 μm i.d.). Peptides were separated with in-house packed column with a 20 cm bed of C18 (Dr. Maisch ReproSil-Pur C18-AQ, 120 Å, 3 μm) having an integrated fritted tip (360 μm o.d.), 75 μm i.d., 15 μm i.d. tip; New Objective). Data-dependent acquisition was performed by selecting top precursor ions for fragmentation using collision-induced dissociation (CID) with 30 sec dynamic exclusion time limit.

#### Orbitrap Fusion Lumos

B31, B31-5A4, and MM1 pooled fractions – 14 fractions per isolate – were analyzed on a Vanquish Neo UHPLC coupled to an Orbitrap Fusion Lumos instrument (Thermo Scientific, USA), equipped with a Easy-Spray nanoelectrospray source. Peptides were loaded onto a trap column (0.5 cm × 300-μm i.d., stationary phase C18) with a flow rate of 10 µL/min of mobile phase: 98% (vol/vol) LC-MS solvent A [0.1% (vol/vol) formic acid (FA) in water] and 2% (vol/vol) LC-MS solvent B [0.1% (vol/vol) FA in acetonitrile]. Peptides were chromatographically separated on a 50-cm analytical column [(EASY-Spray ES803A, Thermo Scientific); 75 µm × 50 cm, PepMap RSLC C18, 2-µm i.d, 100-Å-pore-size particles] applying a 115-min linear gradient: from 3% solvent B to 8% solvent B in 10 min, to 30% solvent B in 90 min, and ramped to 80% solvent B in 5 min, at a flow rate of 250 nL/min. The column temperature was set to 45 °C. Spray voltage was set to 1.8 kV and s-lens RF levels at 30%. The mass spectrometer was set to high resolution data-dependent acquisition (DDA) of 15 topN most intense ions with charge state of +2 to +5. Each MS1 scan (120,000 resolving power at 200 *m/z*, automated gain control (AGC) of 125%, scan range 300 to 1,500 *m/z*, and dynamic exclusion of 30 s, with maximum fill time of 50 ms) was followed by 15 MS2 scans (30,000 resolving power at 200 *m/z*, AGC of 200%, maximum fill time of 54 ms). Higher-energy collisional dissociation (HCD) was used with 1.6 *m/z* isolation window and normalized collision energy of 30%.

#### Triple-TOF

Log phase B31 and MM1 *Borrelia* samples were analyzed using 5600+ Triple-TOF mass spectrometry (ABSciex, USA) coupled with Eksigent 400 nano-HPLC (Sciex, USA). Peptides from the 2 isolates were run separately by loading on trap column (200 μm × 0.5 mm, Chrom XP C18-CL 3 μm, 120 Å, Eksigent, AB Sciex). Peptides were separated on analytical column (75 μm × 20 cm, ChromXP C18-CL 3 μm, 120 Å, Eksigent, Sciex) with the gradient of buffer B (95% acetonitrile in 0.1% formic acid) and flowrate was 300 nL/min. The linear gradient profile from 3 to 40% buffer B in 103 min, increased to 80% in 105 min and continued to 113 min.Buffer B was then brought down to 3% in 115 min and continued until 140 min. Precursor mass was measured at MS1 level in high resolution mode with mass range of 400-1250 *m/z*. The TOF-MS parameters includes: nanospray ionization, curtain gas (CUR)-25, ion source gas 1 (GS1)-3, interface heater temperature (IHF)-150, ion spray voltage floating (ISDF)-2300, declustering potential (DP)-100, collision energy (CE)-10, accumulation time-50 ms, mass tolerance 100 ppm, exclude former peptide ion-15 sec after first detection and precursors selected for each cycle top 30 intense peaks with charge state 2 to 4 having greater than or equal to 150 counts were selected for fragmentation using rolling collision energy. Similarly, at MS2 level, spectra were collected in m/z range of 100-1500 m/z with 50 ms accumulation time in high sensitivity mode.

#### Tims-TOF PRO

All 14 MM1 pooled fractions from high pH fractionation were spiked in with iRT standard peptides (Biognosys AG, Schlieren, Switzerland) and subjected to mass spectrometry (MS) analysis using a timsTOF PRO mass spectrometer (Bruker), coupled to a Vanquish Neo HPLC system (Thermo-Fisher Scientific) in nanoflow setup for both Data-Dependent Acquisition-Parallel Accumulation-Serial Fragmentation (DDA-PASEF) and Data-Independent Acquisition (DIA) PASEF modes. Both modes were operated with 99.9% water, 0.1% formic acid/Milli-Q water (v/v, buffer A), and 99.9% ACN, 0.1% formic acid (v/v, buffer B). Peptides were trapped on a 0.5 cm x 0.3 mm trap cartridge Chrom XP C18, 3 μm (Thermo-Fisher Scientific) at 10 μL/min, and separated on a C18 UHP 15 cm x 0.15 mm × 1.5 μm column (Bruker/PepSep) at either 600 nL/min or 1 μL/min for 66 and 45 minutes, respectively. The gradient elution profile for both flow rates was as follows: 3% to 25% B in 51 min (37 min for 1 μL/min), 25% to 35% B in 15 min (8 min for 1 μL/min), 35% to 80% B in 1 min, followed by an isocratic flow at 80% B for 2 min. The Captive Spray ion source was equipped with a 20 μm emitter (Bruker) and the parameters were as follows: 1700 V Capillary voltage, 3.0 L/min dry gas, and temperature set to 180 °C. The DDA-PASEF data covered 100–1700 m/z range with 6 (for 45 min gradient length) or 8 (for 66 min) PASEF ramps. The TIMS settings were 100 ms ramp and accumulation time (100% duty cycle), resulting in 0.9 s (45 min) 1.1 s (66 min) of total cycle time. Active exclusion was enabled with either a 0.2 (45 min) and 0.3 (66 min) min release. The default collision energy with a base of 0.6 1/K0 [V s/cm^2^] is set at 20 eV and 1.6 1/K0 [V s/cm^2^] at 59 eV was used. Isolation widths were set at 2 m/z at <700 m/z and 3 m/z at >800 m/z. To achieve more comprehensive coverage, 14 fractions were acquired using DIA-PASEF preformed py5 scheme (Bruker) with 32 X 25 Da windows, covering the m/z range of 400-1200 and 1/K0 range of 0.6 to 1.42, resulting in a total cycle time of 1.8 s.

### Proteomic data analysis

In-house developed Trans-Proteomic Pipeline TPP v6.2.0 Nacreous, Build 202302160135-8863 was used for the mass spectrometry data analysis for both identification and quantitation of the proteins. Mass spectrometry raw data (.raw, .d, and .wiff files) from in-house performed experiments and public datasets were converted into .mzML files using msConvert 3.0.5533 [37] and AB_SCIEX_MS_Converter 1.3 Beta from AB SCIEX. The converted files were searched using comet version 2023.01 rev. 0 [38]. All files were searched against a combined reference database, which comprised the following genome assemblies and proteomes. For isolate B31, the Uniprot [39] proteome (ProteomeID UP000001807 [9, 14]), with 1,291 protein sequences (Table 1). This database was named “core proteome” in the build. Also, the RefSeq [40] assembly with accession GCF_000008685.2 containing 1,359 protein sequences, and the GenBank [41] assembly GCA_000008685.2 with 1,339 sequences. Total number of non-redundant protein sequences for isolate B31 is 1,485. For isolate B31-5A4, the GenBank assembly GCA_024662195.1 with 1,429 protein sequences, the RefSeq assembly GCF_024662195.1 with 1,354 sequences, and the ISB assembly with 814 sequences (not published). The total number of non-redundant protein sequences for isolate B31-5A4 is 1,443. For isolate MM1, the GenBank assembly GCA_003367295.1 with 1,302 protein sequences, and the RefSeq assembly GCF_003367295.1 with 1,159 sequences, and an overall total of 1,383 non-redundant protein sequences (Table 1). All public protein databases were downloaded on April 7^th^ 2023. The final combined protein databaseincluded 116 contaminant sequences from cRAP database (http://www.thegpm.org/crap/), downloaded on July 22^nd^ 2022 (Table 1), containing all 3 isolates with 2,619 unique sequences and an equal number of decoy sequences (generated using the decoy tool in Trans-Proteomic Pipeline with “randomize sequences and interleave entries” decoy algorithm) The following data analysis parameters were used: peptide mass tolerance 20 ppm, fragment ions bins tolerance of 0.02 *m/z* and monoisotopic mass offset of 0.0 *m/z for Q-Exactive and Orbitrap Fusion Lumos,* fragment ions bins tolerance of 1.0005 *m/z* and a monoisotopic mass offset of 0.4 *m/z for LTQ Orbitrap Elite/XL,* peptide mass tolerance 20 ppm, fragment ions bins tolerance of 0.1 *m/z* and monoisotopic mass offset of 0.0 *m/z for Triple-TOF and Tims-TOF,* peptide mass tolerance 3.1 Da, fragment ions bins tolerance of 1.0005 *m/z* and monoisotopic mass offset of 0.4 *m/z for LTQ/LCQ Duo/amaZon ion trap, semi-tryptic peptides, allowed 2 missed cleavages, static modification-* carbamidomethylation of cysteine (+57.021464 Da) and variable modifications-oxidation of methionine and tryptophan (+15.994915 Da), protein N-terminal acetylation (+42.0106), peptide N-terminal Gln to pyro-Glu (−17.0265), peptide N-terminal Glu to pyro-Glu (−18.0106), phosphorylation of Ser, Thr, or Tyr (+79.9663). PeptideProphet was used to assign the scores for peptide spectral matches (PSM) for individual files and iProphet was used to assign the score for peptides [<u>31</u>, <u>42</u>, <u>43</u>]. Uniprot proteomes are available at https://www.uniprot.org/proteomes/, and NCBI RefSeq and GenBank genome assemblies are available at https://www.ncbi.nlm.nih.gov/assembly/.

**Table 1.** Number of protein sequences per reference database.

| #proteins | B31 | B31-5A4 | MM1 |
| --- | --- | --- | --- |
| RefSeq | 1,359 | 1,354 | 1,159 |
| GenBank | 1,339 | 1,429 | 1,302 |
| UniProt | 1,291 |  |  |
| ISB |  | 814 |  |
| Total non-redundant | 1,485 | 1,443 | 1,383 |

### PeptideAtlas Assembly

The iProphet outputs from *Q-Exactive, Orbitrap Fusion Lumos*, LTQ Orbitrap Elite/XL, Tims-TOF DDA and *Triple-TOF runs* werefurther processed using two round of reSpect to identify chimeric spectra[44]. For the first round of reSpect, the MINPROB was set to 0 and the MINPROB was set to 0.5 for the second round of reSpect. The new set of .mzML files generated by both rounds of reSpect were searched using comet with the precursor mass tolerance 3.1 and isotope_error off, and processed using the TPP as for the initial files. Using the PeptideAtlas processing pipeline, all the iProphet results from standard and reSpect were filtered at a variable probability threshold to maintain a constant peptide-spectrum match (PSM) FDR of 0.05% for each experiment. The filtered data was assessed with the MAYU software [45] to calculate decoy-based FDRs at the peptide-spectrum match (PSM), distinct peptide, and protein levels. PTMProphet [46] was used to access the localization confidence of the sites with post-translational modifications (PTMs), and for low resolution ITCID runs DALTONTOL=0.6 and DENOISE NIONS=b parameters were applied. Otherwise, default options were used. Bio Tools SeqStats (https://metacpan.org/pod/Bio::Tools::SeqStats) was used to get protein molecular weight, length, pI, and GRAVYin scores [47]. All results were collated in the Borrelia PeptideAtlas, made available at http://www.peptideatlas.org/builds/borrelia/.

### Label-free quantitation

StPeter was used for a label-free quantitation of the build data using spectral counting through a seamless interface in the TPP [48]. The merged protein databases were clustered using OrthoFinder [49]. The representative protein sequence from each protein cluster was extracted. The protein database of the PTMProphet output from each experiment was refreshed to the representative protein database mapping using the RefreshParser tool in TPP. ProteinProphet and StPeter were run on the updated PTMProphet file. The StPeter FDR cutoff value 0.01 and minimum probability 0.9 were used. For FTMS HCD/CID and Tims-TOF runs, a mass tolerance of 0.01 was used.

### RNA transcript analysis

To generate RNA for sequencing, *B. burgdorferi* isolates B31, MM1, and B31-5A4 were cultured as previously described, and the cells were collected by centrifugation. Total RNA was extracted using Qiagen RNEasy Mini kits (Qiagen, USA) according to the manufacturer’s instructions, including an on-column DNAse digestion step. RNA concentration was measured using a NanoDrop spectrophotometer (Thermo-Fisher Scientific, USA) and quality assayed by Agilent BioAnalyzer (Agilent, USA). Prior to library construction, 1 µg of total RNA was depleted of ribosomal-RNA transcripts using MICROBExpress Bacterial mRNA Enrichment Kits (Thermo-Fisher Scientific, USA). Libraries were prepared using NEBNext Ultra II Directional RNA Library Prep Kit for Illumina (New England Biolabs, USA) and NEBNext Multiplex Oligos for Illumina (NEB, USA). The libraries were prepared according to manufacturer’s instructions with insert size approximately 400 bp. Library quality was validated by Agilent Bioanalyzer and yield measured by Qubit HS DNA assay (Thermo Fisher Scientific, USA). Libraries were run on an Illumina NextSeq500 sequencer with High Output Flowcell (Illumina, USA) for 150 cycles. Reads were mapped to the B31 reference genome (Genbank assembly accession GCA_000008685.2) using STAR [50] with quantMode enabled. Mapped reads were visualized with Integrative Genomics Viewer [51] and counts normalized in reads per kilobase of transcript per million reads mapped (RPKM).

## Data Record 1

Mass spectrometry data from 9 public datasets, comprising a total of 617 DDA-MS runs, of isolates B31, B31-ML23, B31-A3, and 297, were used for the Borrelia PeptideAtlas assembly through TPP analysis, with the following identifiers: data on isolate B31 PeptideAtlas dataset PASS00497 [16], ProteomeXchange datasets PXD010065 [17], PXD007904 [21], PXD002365 [22], PXD001860 [23], and MassIVE dataset MSV000085503 [19]; data on isolate B31-ML23 PXD015685 [27], data on isolate B31-A3 PXD005617 [18]; and data on isolate 297 was obtained from the public resource PXD000876 [20] (Supplementary Table S1).

## Data Record 2

Mass spectrometry data from 17 different in-house experiments using laboratory isolates B31, B31-5A4, and MM1, with a total of 210 DDA- and 28 DIA-MS runs (Thermo Scientific instrument .raw files, Bruker instruments .d files), were analyzed through the TPP pipeline and deposited to the ProteomeXchange Consortium via the PRIDE [52] partner repository with the dataset identifier PXD042072.

## Technical Validation

### Borrelia PeptideAtlas assembly

The Borrelia PeptideAtlas repository contains information on peptides identified by mass spectrometry-based proteomics of different infective (B31-5A4, B31-A3, 297) and non-infective (B31, B31-ML23, MM1) *B. burgdorferi* laboratory isolates. The current build (2023-05) comprises extensive proteomics analysis on the total proteome, the secretome and the membrane proteome of the isolates from 9 public datasets and 17 in-house performed experiments with a total of 26 experiments and 855 MS runs. To generate the build, the dense MS-based proteomic data, which includes 57 million MS/MS spectra, was searched using combined reference databases of B31, B31-5A4 and MM1, and uniformly processed through the TPP (see “Methods”). This approach includes the use of the post-search engine reSpect to boost peptide identification from chimeric spectra [44] and MAYU [45] to help estimate decoy-based FDR levels for the Borrelia build, which include multiple large datasets. This strategy allowed the match of approximately 8 million PSMs with FDR level threshold less than 0.0005 at the PSM level, and identification of a total of 76,936 distinct peptides at 0.1% peptide FDR (Figure 3a). These peptides mapped to a total of 1,581 proteins among all isolates with a protein-level FDR less than 1% (Figure 3b), including 924 core canonical and 297 noncore canonical. The description of all protein categories and a summary of the proteins identified within each category in the build are shown in Table 2, and complete information on proteins identified in the build is made available in Supplementary Table S2. Specifically, for the B31 core proteome, 1,107 non-redundant proteins to which at least one peptide was mapped were identified, covering 86% of the B31 core proteome (Table 3). Figure 3 c-e shows the frequency distributions of observed and theoretical tryptic peptides by length (aa), distributions of peptide charge and the number of distinct peptides per million observed in each isolate experiment, respectively. The majority of the identified peptides had a charge state of 2+ or 3+ with a length of 7 to 30 amino acids, and most of the identified peptides presented at least one trypsin missed cleavage site. Figure 3f illustrates the frequency (%) of the primary sequence coverage for canonical proteins, i.e., the percentage value of amino acids which were identified for each protein, which ranged from 6% to 100%. The complex and detailed proteomic results achieved with the Borrelia PeptideAtlas repository were made available at http://www.peptideatlas.org/builds/borrelia/.

**Figure 2.**
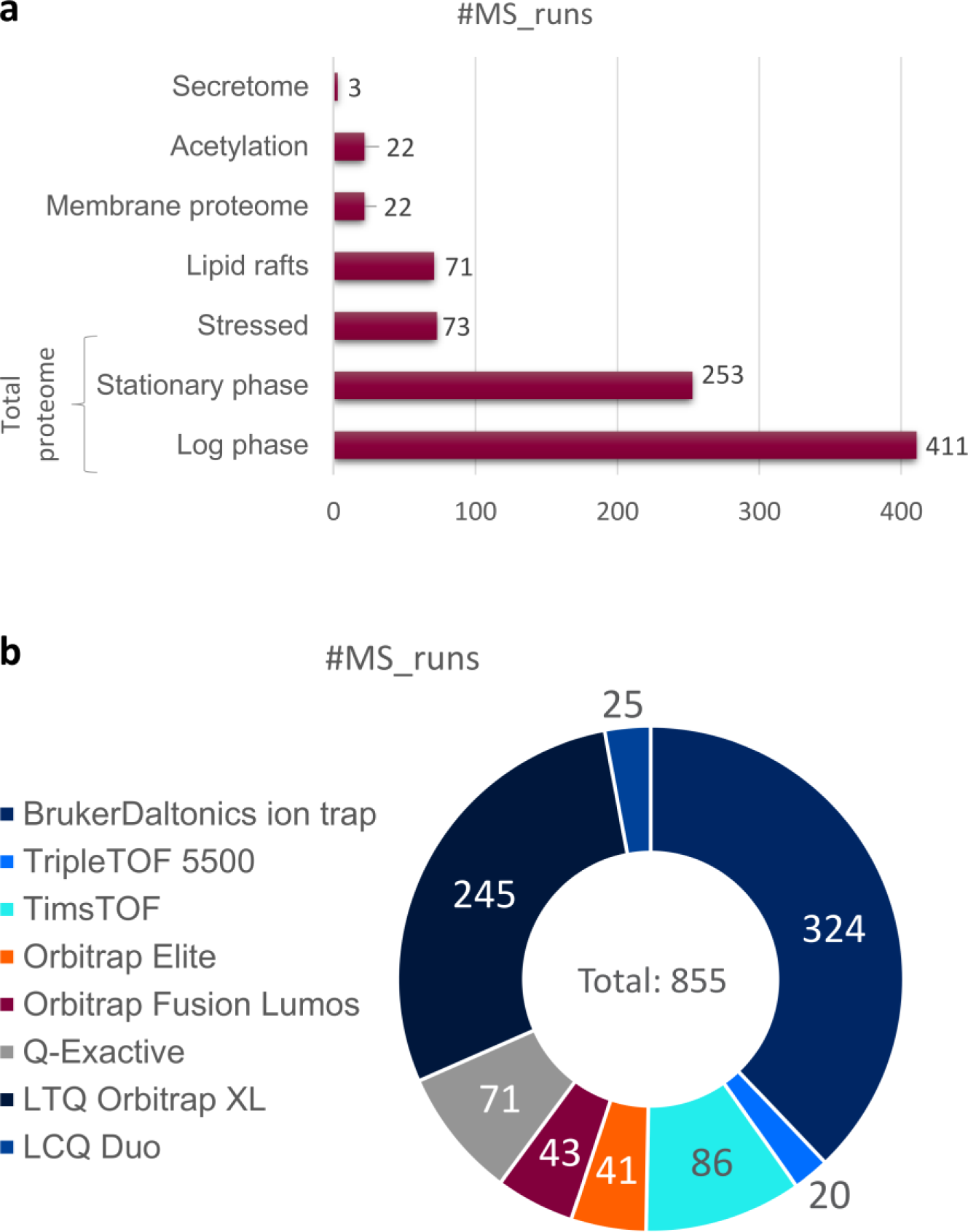
Mass spectrometry runs. (a) Number of MS runs per environmental condition of *B. burgdorferi* analyzed for total proteome, or protein enrichment methodology (secretome, acetylation, membrane proteome, lipid rafts). (b) Number of MS runs per instrument used. Bruker instruments: BrukerDaltonics ion trap, TripleTOF 5500, TimsTOF. Thermo Scientific instruments: Orbitrap Elite, Orbitrap Fusion Lumos, Q-Exactive HF, LTQ Orbitrap XL, LCQ Duo.

**Figure 3.**
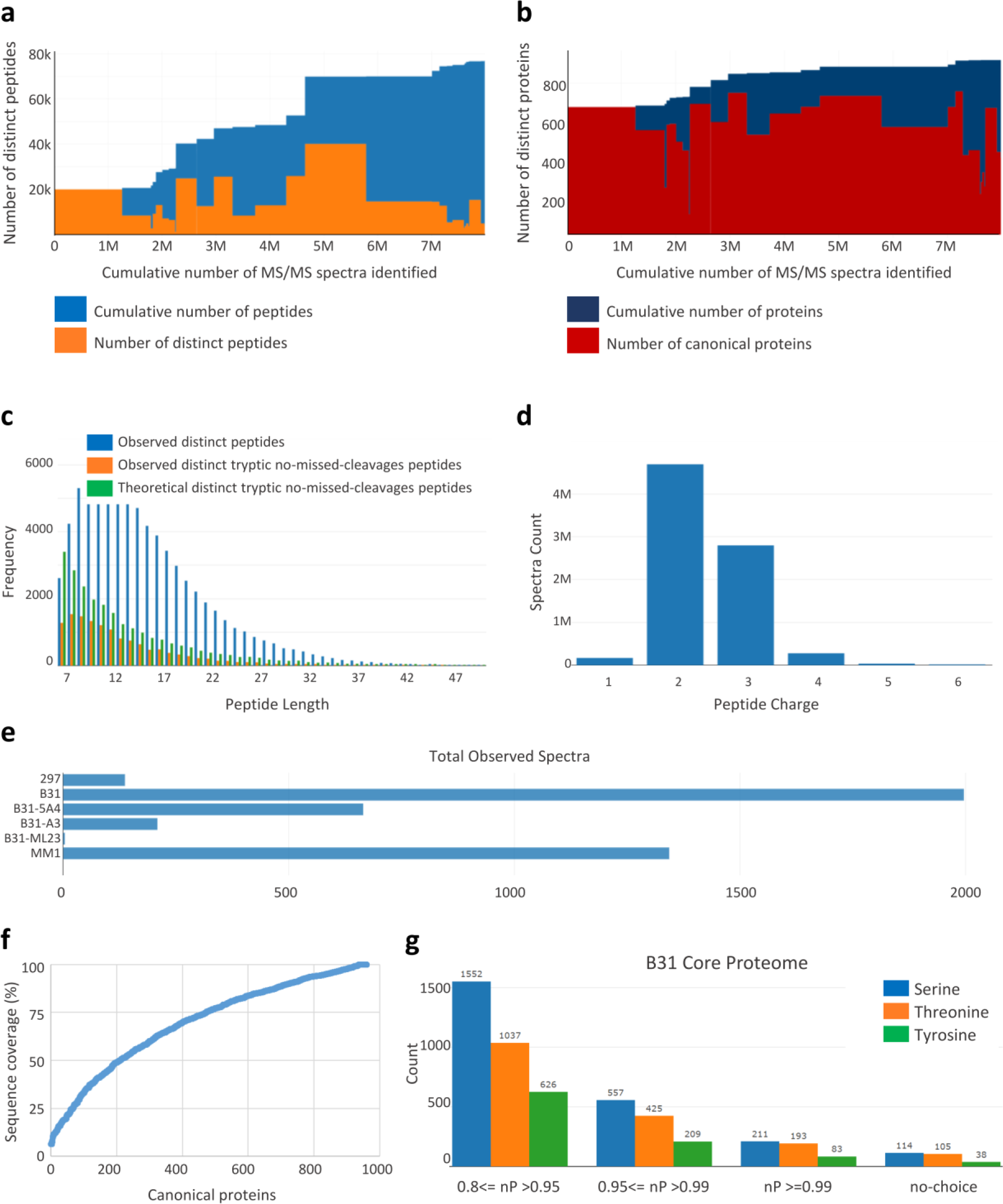
Borrelia PeptideAtlas experiment contribution. (a) Number of peptides which contributed to each experiment, and the cumulative number of distinct peptides for the build as of that experiment. (b) Cumulative number of canonical proteins contributed by each experiment. Height of red bar is the number of proteins identified in experiment; height of blue bar is the cumulative number of proteins; width of the bar (x-axis) shows the number of spectra identified (PSMs), above the threshold, for each experiment. (c) Frequency distributions of peptide length by number of amino acids. The figure shows frequency of distinct peptides (in blue), distinct tryptic peptides with no missed cleavages (in orange), and theoretical, i.e., not observed, tryptic peptides with no missed cleavage (in green). (d) Frequency distributions of peptide charge. (e) Number of distinct peptides per million observed in each isolate experiments. (f) Relative protein sequence coverage for canonical proteins based on sequence coverage, i.e., the % of amino acids of the primary sequence which were identified. (g) Histogram showing the frequency distribution of PSMs of phosphorylated sites (serine, threonine, and tyrosine), identified for B31 UniProt core proteome, according to PTMProphet probability (nP). nP ranges from 0.8 to 0.99. no-choice: shows PSMs with only one possible phosphorylation site available, hence nP=1. Blue, yellow, and green bars indicate serine, threonine, or tyrosine phosphorylated sites, respectively.

**Table 2.**
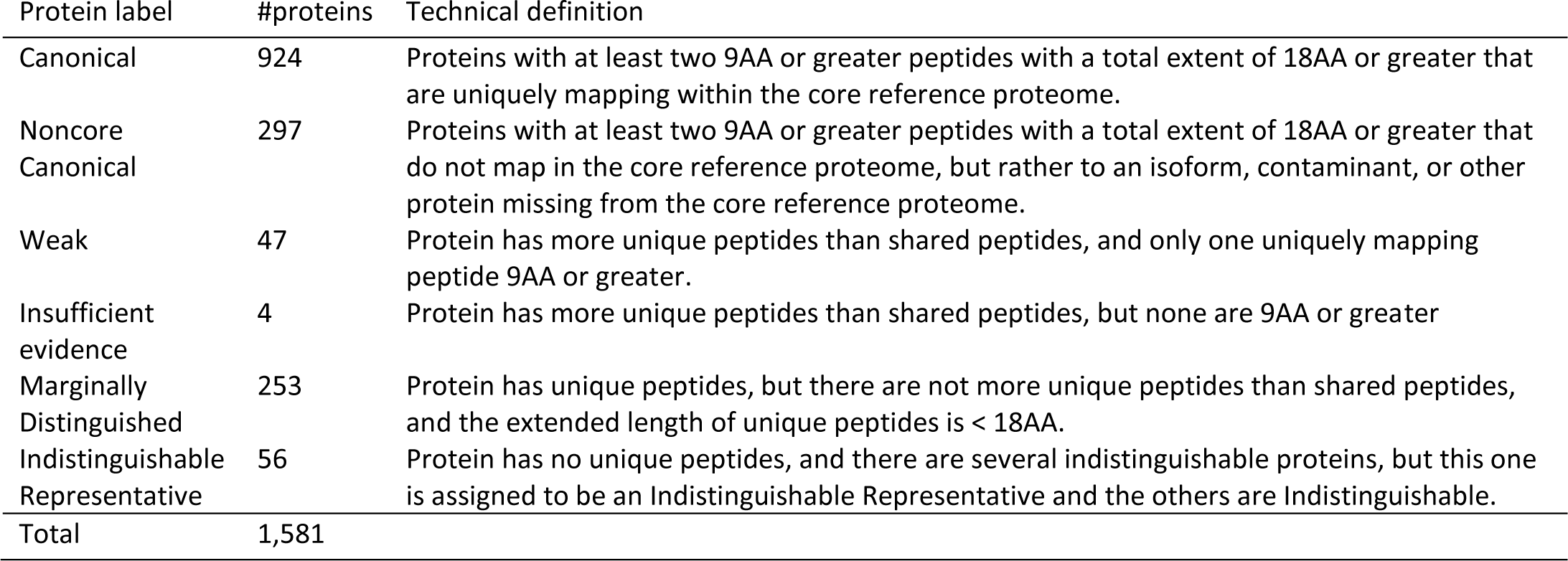
Protein identification categories in the Borrelia PeptideAtlas build.

| Protein label | #proteins | Technical definition |
| --- | --- | --- |
| Canonical | 924 | Proteins with at least two 9AA or greater peptides with a total extent of 18AA or greater that are uniquely mapping within the core reference proteome. |
| Noncore Canonical | 297 | Proteins with at least two 9AA or greater peptides with a total extent of 18AA or greater that do not map in the core reference proteome, but rather to an isoform, contaminant, or other protein missing from the core reference proteome. |
| Weak | 47 | Protein has more unique peptides than shared peptides, and only one uniquely mapping peptide 9AA or greater. |
| Insufficient evidence | 4 | Protein has more unique peptides than shared peptides, but none are 9AA or greater |
| Marginally Distinguished | 253 | Protein has unique peptides, but there are not more unique peptides than shared peptides, and the extended length of unique peptides is < 18AA. |
| Indistinguishable Representative | 56 | Protein has no unique peptides, and there are several indistinguishable proteins, but this one is assigned to be an Indistinguishable Representative and the others are Indistinguishable. |
| Total | 1,581 |  |

**Table 3.** Proteome coverage. Database: name of database, which collectively form the reference database for this build. #entries: total number of entries. #proteins: total number of non-redundant entries. #obs-proteins: number of non-redundant protein sequences within the subject database to which at least one observed peptide maps. %observed: the percentage of the subject proteome covered by one or more observed peptides. #unObs-proteins: number of non-redundant protein sequences within the subject database to which no observed peptide maps.

| Database | #entries | #proteins | #obs-proteins | %observed | #unObs-proteins |
| --- | --- | --- | --- | --- | --- |
| B31 CoreProteome | 1,291 | 1,291 | 1,107 | 85.7 | 184 |
| B31 | 3,989 | 1,485 | 1,235 | 83.2 | 250 |
| MM1 | 2,461 | 1,383 | 1,159 | 83.8 | 224 |
| B31-5A4 | 3,597 | 1,443 | 1,221 | 84.6 | 222 |

### Post-translational modifications

For protein phosphorylation analysis of the Borrelia PeptideAtlas each dataset was further analyzed by PTMProphet, embedded in the TPP pipeline, to compute localization probabilities (*P*) of phosphor-sites, including: serine (pS), threonine (pT) and tyrosine (pY) residues (Figure 3g). PTMProphet applies Bayesian models for each passing PSM that contains a phosphor PTM as reported by the search engine [46]. PTMProphet probabilities for STY-sites present in the Borrelia build range from 0 to 1 (highest significance), with greater values indicating higher probability that a phosphate group is present at the site, based on MS/MS evidence [46]. The complete information on PTMProphet analysis for all 4 databases (B31 core proteome, B31, B31-5A4, and MM1) is made available in Supplementary Table S3. Specifically, in the B31 core proteome, the total number of potential phosphor sites among the observed proteins is 25,547 for serine, 14,296 for threonine and 14,788 for tyrosine. The number of potential phosphorsites with peptide coverage among these proteins is 2,711 (10.61%) for serine, 1,720 (12.03%) for threonine and 1,153 (7.80%) for tyrosine. Among these a total of 211 phospho-serine sites, 193 phospho-threonine sites and 83 phospho-tyrosine sites were identified with PTMProphet probability ≥0.99. Considering all phosphor-sites (STY) with *P* ≥ 0.99 identified in all canonical proteins in the build, including the redundancy of phosphor-sites, a total of 42,156 phosphor-sites were seen throughout 1,542 proteins (Supplementary Table S3).

During its life *B. burgdorferi* is exposed to different environmental conditions while cycling through ticks and mammalian hosts, including: changes in temperature, pH and nutrient sources [53]. Furthermore, *B. burgdorferi* lacks genes of the tricarboxylic acid cycle and oxidative phosphorylation and is not capable of *de novo* biosynthesis of carbohydrates, amino acids, or lipids; instead relying on the host’s metabolism. Protein phosphorylation in *B. burgdorferi* has been described with a critical role in the pathogen’s growth and chemotaxis signal transduction [54]. During the tick phase, specifically, *B. burgdorferi* relies uniquely on glycolysis for ATP production [53]. Glycerol is a carbohydrate readily available in ticks, and once transported to the spirochete cytoplasm it is phosphorylated by the glycerol kinase GlpK to generate glycerol 3-phosphate, which will follow the glycolytic cascade [55]. Here, we use the glycerol kinase GlpK as an example of a phosphorylated protein to show the Borrelia PeptideAtlas interface (Figure 4). GlpK is a canonical protein identified with 171 phospho-sites with *P* ≥ 0.99. Figure 4 shows the Borrelia PeptideAtlas interface after searching results for GlpK protein identifier (UniProt entry O51257) in the build protein browser. Figure 4a displays the GlpK primary sequence coverage of 100%, and Figure 4b illustrates the distribution of all observed distinct peptides for that protein. It is possible to open the peptide browser for each peptide by clicking on the individual blue bar. In the same page, it is possible to visualize pSTY-sites distributed in the protein sequence, with the corresponding PTMProphet probabilities (Figure 4c), and a view table with information on the distinct observed peptides, which contain the phospho-sites (Figure 4d). The Borrelia PeptideAtlas PTM summary can be accessed at http://www.peptideatlas.org/builds/borrelia/, in the “PTM coverage” section.

**Figure 4.**
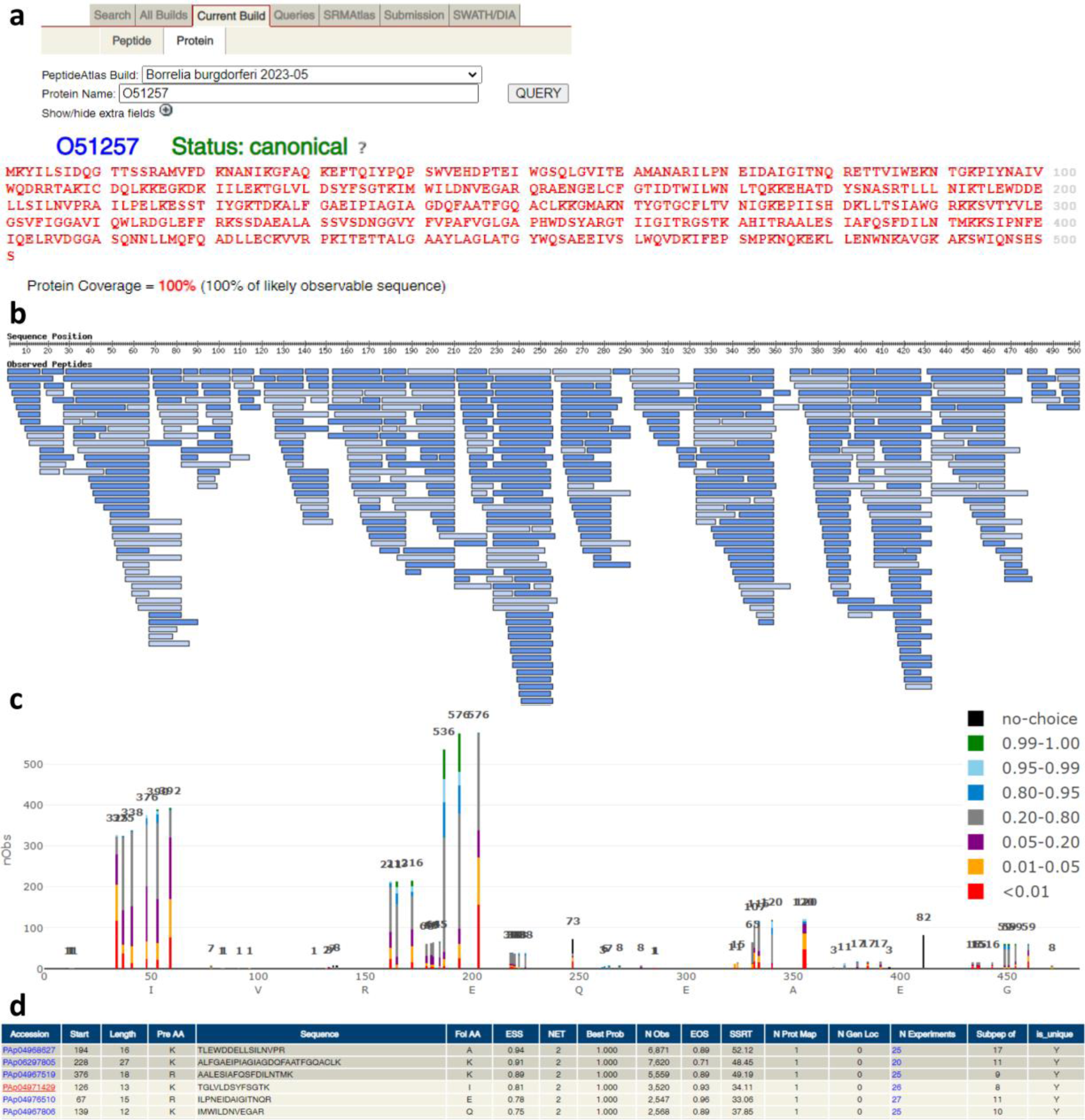
Borrelia PeptideAtlas view of glycerol kinase (gene name GlpK, UniProt entry O51257) phosphorylated sites. Example of the protein PTM summary on the Borrelia PeptideAtlas. (a) View of the protein search tab and corresponding primary protein sequence coverage, in red. (b) View of the primary protein sequence display with observed peptides. (c) Distribution of phosphorylated sites in OspC protein sequence with PTMProphet probabilities, ranging from less than 0.01 to 1. (d) Information on observed peptides including empirical suitability score (ESS) empirical observability score (EOS). Accession: peptide accession; start: start position in the protein; pre AA: preceding (towards the N terminus) amino acid; sequence: amino acid sequence of detected peptide, including any mass modifications; fol AA: following (towards the C terminus) amino acid; ESS: empirical suitability score, derived from peptide probability, EOS, and the number of times observed. This is then adjusted sequence characteristics such as missed cleavage [MC] or enzyme termini [ET], or multiple genome locations [MGL]; NET: highest number of enzymatic termini for this protein; NMC: lowest number of missed cleavage for this protein; Best Prob: highest iProphet probability for this observed sequence; Best Adj Prob: highest iProphet-adjusted probability for this observed sequence; N Obs: total number of observations in all modified forms and charge states; EOS: empirical Observability Score, a measure of how many samples a particular peptide is seen in relative to other peptides from the same protein; SSRT: Sequence Specific Retention time provides a hydrophobicity measure for each peptide using the algorithm of Krohkin et al. Version 3.0 [56]; N Prot Map: number of proteins in the reference database to which this peptide maps; N Gen Loc: number of discrete genome locations which encode this amino acid sequence; Subpep of: number of observed peptides of which this peptide is a subsequence.

### Genome coverage of *B. burgdorferi* isolates

Due to the variability of the plasmid content in different *B. burgdorferi* isolates – which account for approximately one-third of the genome [57], combined reference databases of laboratory isolates B31, B31-5A4 and MM1 were used to search the dense proteomic data when constructing the build. These databases comprise reference genome assemblies from NCBI RefSeq, GenBank, and UniProt proteome (see “Methods”). As aforementioned, isolate B31 genome contains a linear chromosome (843 genes) and 21 plasmids (12 linear and 9 circular, 670 genes and 167 pseudogenes total) [14]. Of the 1,513 genes, 1,291 are predicted as unique protein-coding genes. The infective B31-5A4 genome assembly indicates the presence of, besides the linear chromosome, 11 linear plasmids and 9 circular plasmids (ISB, not published). Isolate MM1 has 15 plasmids (7 linear and 8 circular), including the unique lp28-8 and the conserved chromosome [58].

The linear chromosome carries approximately 65% of all genes in *B. burgdorferi*, which encode housekeeping proteins involved in DNA replication, transcription and translation regulation, besides energy metabolism [14]. Here, more than 95% of proteins encoded by the chromosome genome were identified with FDR levels less than 1% throughout all isolates (Figure 5; Supplementary Table S2). Circular plasmid cp26 and linear plasmid lp54 are stable and present in all *B. burgdorferi* isolates studied to date [59], including B31, B31-5A4 and MM1, and hence considered a control for encoded proteins identified in the build (Figure 6). Plasmid cp26 encodes proteins which are essential for early stages of infection in mammalian hosts, e.g. outer surface protein C (OspC) [60]. Thus, it is considered an essential plasmid for the spirochete growth and survival [41]. Similarly to cp26, the linear plasmid lp54 is present in all *B. burgdorferi* genotypes and encodes critical proteins in tick colonization, e.g. surface proteins OspA and OspB, in tissue attachment and proliferation, such as Decorin-binding proteins A and B, and Crasp1, which plays a critical role in evasion of the host immune system by binding proteins of the complement system [61]. Accordingly, 96% of proteins encoded by cp26 had peptide coverage for B31, 100% for B31-5A4, and 93% for MM1; and around 85% of proteins encoded by lp54 had peptide coverage for the 3 isolates (Figure 5; Supplementary Table S2); the remaining plasmids display varying frequencies of proteins identified throughout the isolates, ranging from 37% to 85%. The complete information on non-detected proteins by LC-MS (“missing proteins”) for each isolate reference database is made available in Supplementary Table S4, which includes the plasmid information. We note that 80% of missing proteins are described as hypothetical proteins or of unknown function in UniProt B31 core proteome, 5% are membrane proteins, and the remaining 15% have variable descriptions, including flagellar and transporter proteins.

**Figure 5.**
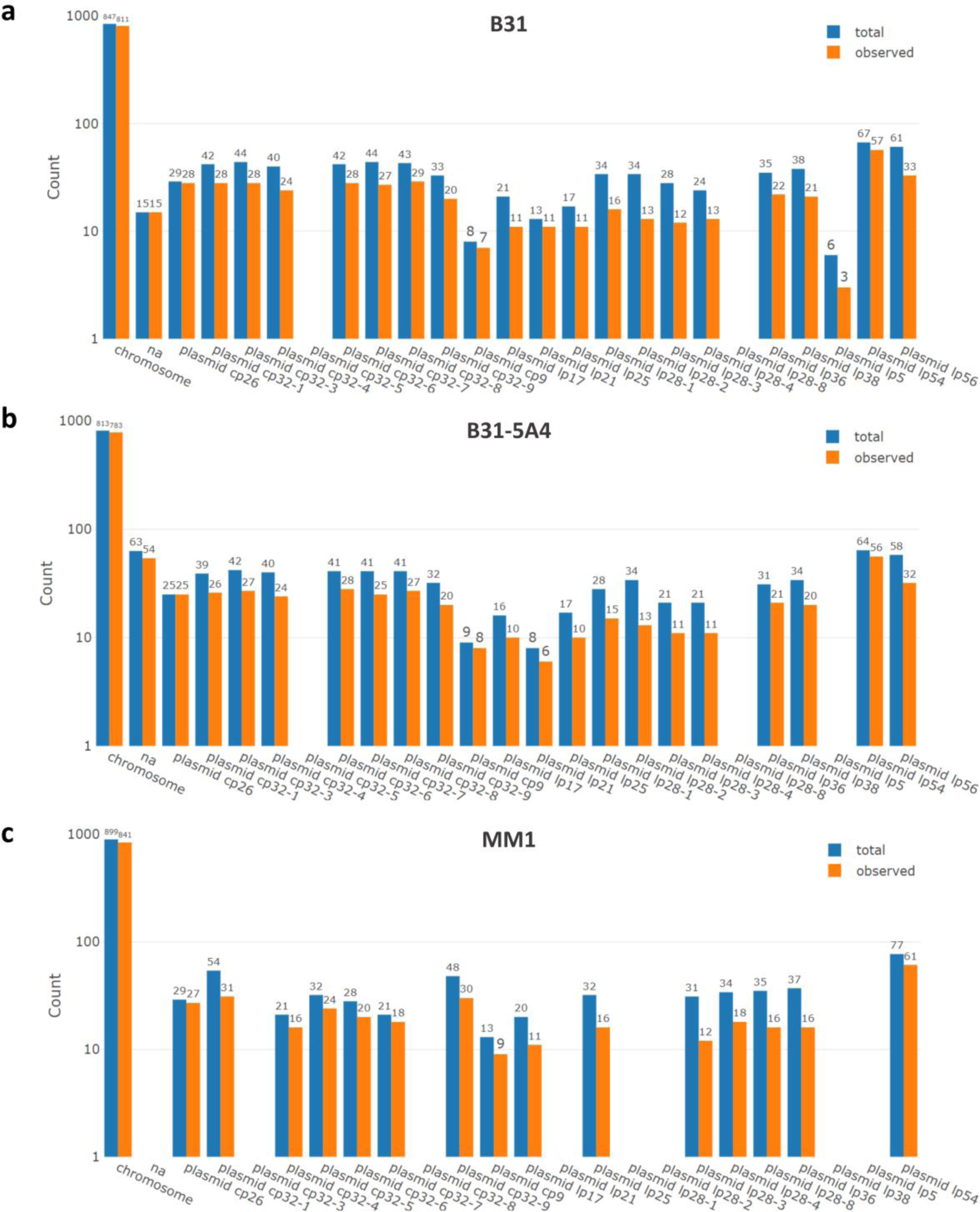
Genome coverage for isolates. Histograms showing the distribution of chromosomal and plasmid coverage for the reference database of isolates B31, B31-5A4, and MM1. Blue bars indicate total number of genes expected for the chromosome or corresponding plasmid. Orange bars indicate number of genes, which correspond to proteins, observed in the chromosome or corresponding plasmid. na: not assigned.

**Figure 6.**
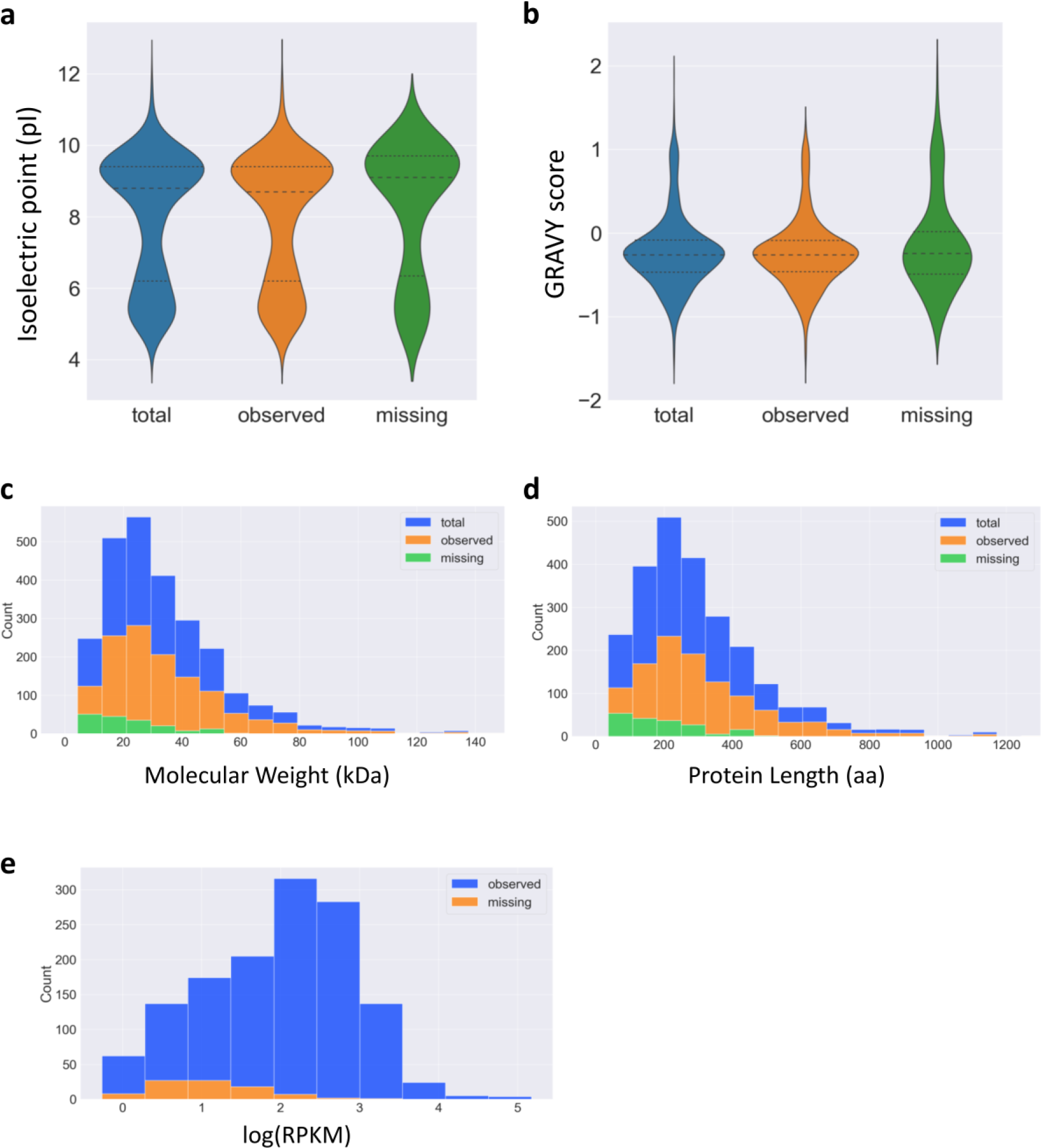
Protein physicochemical properties. Total: number of total proteins in the B31 UniProt reference database (core proteome). Observed: number of observed proteins in the B31 core proteome. Missing: number of proteins not observed in the B31 core proteome. (a,b) Frequency distributions for protein isoelectric point (pI) and GRAVY score, shown as violin plot. Protein GRAVY index score indicates average hydrophobicity and hydrophilicity. GRAVY score below 0 indicates hydrophilic protein, while scores above 0, hydrophobic [47]. (c,d) Frequency distribution for protein molecular weight (kDa) and protein length (number of amino acids), shown as stacked histograms. (e) Frequency distribution of mRNA log_10_ RPKM for observed and not observed (missing) proteins in blue and orange, respectively, shown as a histogram.

Figure 6 shows the physicochemical characteristics of proteins of the B31 core proteome, including total expected proteins in the proteome, observed and missing proteins in the Borrelia PeptideAtlas. The features comprise protein isoelectric point (pI), GRAVY index score, molecular weight (kDa), and length (number of amino acids). The frequency distributions of these features indicate that missing proteins have similar characteristics as those of the observed proteins, with relatively higher frequencies of basic (pI > 10), hydrophobic (GRAVY score > 0) and small proteins (less than 20 kDa) (Figure 6a-d). To further investigate the mRNA levels of the non-detected proteins, transcriptomic analysis of isolates B31, MM1 and B31-5A4 was performed (Supplementary Table S4). Transcripts were not detected by RNAseq for approximately 50% of the missing proteins (Supplementary Table S5). The other 50% have RPKM ranging from 1 to 1,379. A considerable number of canonical proteins detected for the B31 core proteome (around 42%) had low levels of mRNA RPKM, i.e., lower than 100 counts, and the remaining transcripts showed a range of 101-149,599 RPKM (Supplementary Table S4). Therefore, proteins not detected for the B31 core proteome show absence or relatively lower abundance of their corresponding transcripts. The frequency distribution of log_10_ RPKM for transcripts of observed and missing proteins is shown in Figure 6e.

## Usage Notes

The Borrelia PeptideAtlas provides a publicly accessible resource, important for the Lyme disease research community. Our goal is to provide an expandable data source for many other *B. burgdorferi* isolates, including clinically relevant isolates, and subjected to different growth conditions, enabling the investigation of the dynamic proteome of this spirochete through its infection stages and their vastly different environments. The diverse proteomic information from multiple infective isolates with credible data presented by the Borrelia PeptideAtlas can be useful to pinpoint potential protein targets which are common to infective isolates and may be key in the infection process – such as outer membrane proteins. A list of membrane protein targets present in the build can be identified. With *in silico* prediction of signal peptides and secondary structures of membrane proteins, this dense proteomic data can be further investigated for host-pathogen protein interactomics with different technologies. Moreover, this resource provides access to information regarding a wide range of potential proteins and PTMs relevant to chronic, acute and post treatment Lyme disease to develop sensitive diagnostic assays in the Lyme community. The Borrelia PeptideAtlas is a dynamic proteome resource in terms of size and complexity and will be updated to include new data periodically as more genomic and proteomic data is made available for new clinical and laboratory isolates. The collection of the raw data, protein, and peptide information are publically available in the Borrelia PeptideAtlas at http://www.peptideatlas.org/builds/borrelia/.

## Code Availability

The authors do not have code specific to this work to disclose.

## Acknowledgements

This work was funded in part by the National Institutes of Health grants from the National Institutes of Health, National Institute of Allergy and Infectious Diseases (NIAID) R21AI142302 (RLM), R21AI133335 (RLM), and R01AI029735 (MJC), National Institute for General Medical Sciences (NIGMS) R01GM087221, the Office of the Director S10OD026936, and the National Science Foundation award 1920268. We thank the Wilke Family Foundation, and the Steven and Alexandra Cohen Foundation for their generous support.

## Author contributions

Panga J. Reddy: performed *B. burgdorferi* culturing, mass spectrometry data generation experiments and data analysis, PeptideAtlas generation.

Zhi Sun: performed mass spectrometry data analysis, PeptideAtlas generation, and edited the manuscript.

Helisa H. Wippel: performed *B. burgdorferi* culturing, mass spectrometry data generation experiments and data analysis, PeptideAtlas generation, manuscript writing and preparation.

David H. Baxter: performed *B. burgdorferi* genome and RNA sequencing data generation and sequence data analysis.

Kristian E. Swearingen: performed sample preparation experiments. David D. Shteynberg: performed mass spectrometry data analysis. Mukul Midha: performed mass spectrometry experiments.

Melissa J. Caimano: Provided B31-5A4, biological, and technical expertise for preparing *B. burgdorferi* and edited the manuscript.

Klemen Strle: provided Clinical, biological, and technical expertise for preparing *B. burgdorferi* and edited the manuscript.

Yongwook Choi: performed *B. burgdorferi* genome sequence data analysis.

Agnes P. Chan: performed *B. burgdorferi* genome sequence data analysis.

Nicholas J. Schork: performed *B. burgdorferi* genome sequence data analysis and edited the manuscript.

Robert L. Moritz: conceived the project, secured funding, designed experiments, provided technical expertise, managed the project, and performed manuscript writing, preparation, and finalization.

## Competing interests

The authors declare no competing interests.

## Supplementary Table Legends

**Supplementary Table S1. Experiment contribution.** Complete information on public datasets and in-house (ISB) performed experiments. Interactive table is made available at http://www.peptideatlas.org/builds/borrelia/.

**Supplementary Table S2. Borrelia PeptideAtlas identified proteins.** Complete information on proteins identified for the Borrelia build, with FDR levels less than 1%. Description of each PeptideAtlas protein category is included in the table.

**Supplementary Table S3. Phosphorylated sites.** Information on phosphorylated sites – serine, threonine, and tyrosine – covered for the 4 different reference databases: B31 Core Proteome (UniProt), B31, MM1, and B31-5A4, further described in Methods. Description of each category is included in the table.

**Supplementary Table S4. Transcriptomic information.** RNAseq data collected for isolates B31, MM1, and B31-5A4. Strand 2_RPKM: second strand cDNA counts normalized in reads per kilobase of transcript per million reads mapped (RPKM).

**Supplementary Table S5. Information on missing proteins.** Complete information on missing proteins (not observed) in the reference databases of B31, B31-5A4, and MM1.

## Notes

http://www.ebi.ac.uk/pride

http://www.peptideatlas.org

### Competing Interest Statement

The authors have declared no competing interest.

